# The Cell-Cycle Transcriptional Network Generates and Transmits a Pulse of Transcription Once Each Cell Cycle

**DOI:** 10.1101/190686

**Authors:** Chun-Yi Cho, Christina M. Kelliher, Steven B. Haase

## Abstract

Multiple studies have suggested the critical roles of cyclin-dependent kinases (CDKs) as well as a transcription factor (TF) network in generating the robust cell-cycle transcriptional program. However, the precise mechanisms by which these components function together in the gene regulatory network remain unclear. Here we show that the TF network can generate and transmit a “pulse” of transcription independently of CDK oscillations. The premature firing of the transcriptional pulse is prevented by early G1 inhibitors, including transcriptional corepressors and the E3 ubiquitin ligase complex APC^Cdh1^. We demonstrate that G1 cyclin-CDKs facilitate the activation and accumulation of TF proteins in S/G2/M phases through inhibiting G1 transcriptional corepressors (Whi5 and Stb1) and APC^Cdh1^, thereby promoting the initiation and propagation of the pulse by the TF network. These findings suggest a unique oscillatory mechanism in which global phase-specific transcription emerges from a pulse-generating network that fires once-and-only-once at the start of the cycle.

## INTRODUCTION

Genome-wide phase-specific transcription during the cell cycle has been observed in multiple species (Cho et al., 2001; Menges et al., 2003; Rustici et al., 2004; Spellman et al., 1998; Whitfield et al., 2002), yet how this cell-cycle transcriptional program is generated remains poorly understood. Although the biochemical oscillation of cyclin-dependent kinase (CDK) and anaphase-promoting complex (APC) activity has been proposed as the central cell-cycle oscillator that controls phase-specific transcription (Banyai et al., 2016; Rahi et al., 2016), much of the periodic transcriptional program still persists in budding yeast mutant cells whose S-phase and mitotic cyclin-CDK activity are held at constitutively low or high levels (Bristow et al., 2014; Cho et al., 2017; Orlando et al., 2008). By integrating transcriptome analyses and transcription factor (TF) binding localization studies, models have been proposed in which a highly interconnected network of TFs can generate phase-specific transcription via serial activation of transcriptional activators, (Bristow et al., 2014; Lee et al., 2002; Orlando et al., 2008; Pramila et al., 2006; Simmons Kovacs et al., 2012; Simon et al., 2001). However, it is still unclear how the dynamical behaviors of the TF network are feedback-regulated by CDK and APC activities, whose oscillations are also modulated by a complex biochemical network (Chen et al., 2004; Cross, 2003).

The potential of a TF network to oscillate semi-autonomously from CDKs and cell-cycle progression and to trigger cell-cycle transcription has been supported by both Boolean and ODE models (Hillenbrand et al., 2016; Orlando et al., 2008; Simmons Kovacs et al., 2012). However, previous data have also suggested that the amplitude and robustness of cell-cycle transcription are dependent on the presence of CDK activities, particularly those of G1 cyclin-CDKs. In the absence of all Cdc28/Cdk1 activity, global transcript dynamics are severely impaired in arrested G1 cells (Rahi et al., 2016; Simmons Kovacs et al., 2012). On the other hand, in cells expressing G1 cyclins at high levels but lacking S-phase and mitotic cyclins, global cell-cycle transcription persists with dynamics highly similar to that in wild-type cells (Cho et al., 2017; Orlando et al., 2008). Thus, G1 cyclin-CDKs and the TF network function together and are sufficient to trigger a large program of phase-specific transcription. The precise mechanisms by which CDKs promote the robust oscillations of the TF network have not been established.

In mammalian cells, G1 cyclin-CDKs activate G1/S transcription by phosphorylating the transcriptional corepressor Rb and releasing it from the transcriptional activators E2F1-3 (Giacinti and Giordano, 2006). The topology of this network motif is highly conserved in budding yeast (Cross et al., 2011; Johnson and Skotheim, 2013). At Start, Cln3-CDK phosphorylates and inhibits the Rb analogues, Whi5 and Stb1, to relieve their repression on the E2F analogues SBF and MBF (Costanzo et al., 2004; de Bruin et al., 2004; Takahata et al., 2009; Wang et al., 2009). Subsequently, Cln1/2-CDKs are activated and mediate positive feedback loops to fully inhibit Whi5, leading to the coherent G1/S transcription driven by SBF/MBF and the commitment to the cell cycle (Eser et al., 2011; Skotheim et al., 2008).

In this study, we began by asking whether the low-amplitude oscillations observed in the Cdk1 mutant cells (*cdc28-4*) resulted from inefficient inactivation of Whi5/Stb1. Unexpectedly, deletions of *WHI5* and *STB1* genes in a *cdc28-4* background only resulted in constitutively high transcript levels of G1/S genes and low levels of S/G2/M genes. We found that in the absence of Cdk1 activities, APC^Cdh1^ was not fully inactivated, and thus several network TFs were constitutively unstable. Further introduction of a mutation in the gene encoding APC component, Cdc16 (*cdc16-123*), restored the protein levels of network TFs as well as global dynamics of phase-specific transcription. Taken together, our findings suggest that TFs, CDKs, and APC interact in a gene regulatory network to generate and transmit a pulse of transcription as cells progress through the cell cycle. Multiple inhibitory mechanisms, including transcriptional corepressors Whi5/Stb1 in early G1 and repressors in S/G2/M phases, likely restrict the firing of transcriptional pulses to once-and-only-once per normal cell cycle. We propose a unique oscillatory mechanism in which the network produces and propagates a single pulse of transcription upon commitment to the cycle.

## RESULTS

### G1 cyclin-CDKs enhance the generation and transmission of a transcriptional pulse

We sought to determine how G1 cyclin-CDKs (Cln-CDKs) contribute to the generation of the cell-cycle transcriptional program in budding yeast. It has been shown that Cln-CDKs can inhibit the transcriptional corepressors Whi5 and Stb1, which in turn inhibit the transcriptional activating complexes SBF and MBF (Figure 1A) (Costanzo et al., 2004; de Bruin et al., 2004; Skotheim et al., 2008; Takahata et al., 2009; Wang et al., 2009). Once SBF/MBF are derepressed, they activate G1/S transcription of ~200 genes that includes several other transcription factors (Ferrezuelo et al., 2010; Horak et al., 2002). Once transcriptionally activated by SBF/MBF, the partially redundant transcriptional repressors Nrm1/Yhp1/Yox1 mediate negative feedback to attenuate both early-G1 and G1/S transcription and create a transcriptional “pulse” (de Bruin et al., 2006; Pramila et al., 2002). Downstream transcriptional activators, including Hcm1/Plm2/Tos4, SFF, and Swi5/Ace2, are then thought to transmit the G1/S transcriptional “pulse” and sequentially activate transcription in S phase, M phase, and at M/G1 transition (Figure 1A) (Orlando et al., 2008).

**Figure 1.**
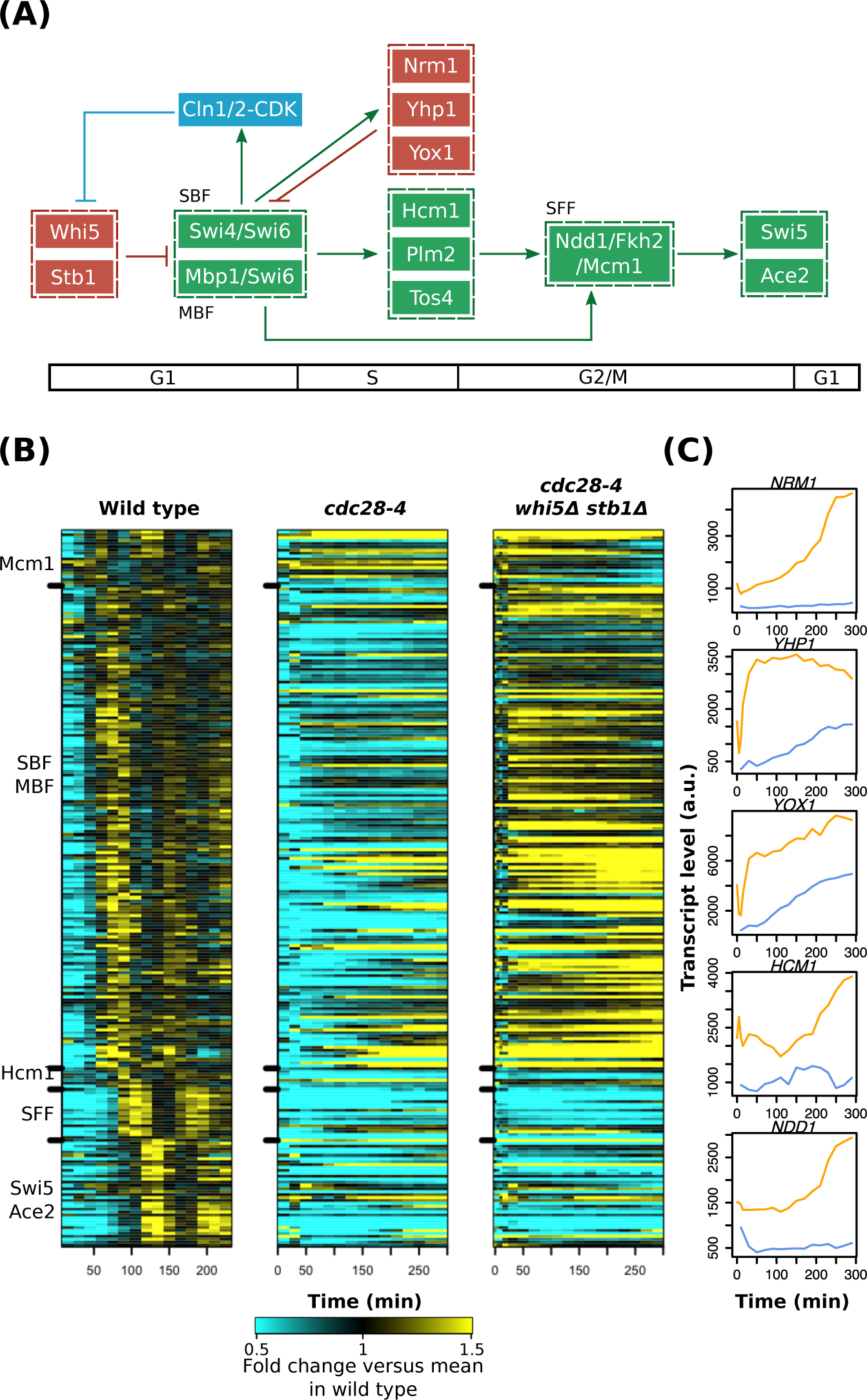
G1 cyclin-CDKs enhance the amplitude of global cell-cycle transcription through inhibiting Whi5/Stb1 and additional mechanisms. (A) A network model for interactions between G1 cyclin-CDKs (blue) and relevant TFs in the network. Transcriptional activators and repressors are shown in green and red, respectively. Nodes are ordered horizontally by their approximate time of activation during the cell cycle. (B) Heat maps depicting transcript dynamics of the canonical genes regulated by the TF network (Table S1) in indicated time courses. Early G1 cells were obtained by centrifugal elutriation and released into YEP-dextrose medium at 37°C. Transcript levels were measured by microarray. Normally cycling wild-type cells from a previous study are shown (Simmons Kovacs et al., 2012). Transcript levels are depicted as fold change relative to mean in wild type. Mcm1 targets are activated in early G1 and are repressed by repressors Yhp1/Yox1 (Pramila et al., 2002); these regulations are not shown in (A) for simplicity of the diagram. (C) Line graphs showing absolute transcript levels of the network TFs in the *cdc28-4/cdk1* mutant (blue) and the *cdk1 whi5Δ stb1Δ* mutant (yellow). See also Figure S1.

In the temperature-sensitive *cdc28-4* mutant cells arrested in G1, only low-amplitude oscillations were observed in a subset of transcripts (Simmons Kovacs et al., 2012). We thus asked whether deletions of the *WHI5* and *STB1* genes in the *cdc28-4* mutant background would restore the dynamics of global cell-cycle transcription. A synchronous G1 population of *cdc28-4 whi5Δ stb1Δ* (denoted as *cdk1 whi5Δ stb1Δ* below) mutant cells were collected by centrifugal elutriation and then released into YEP-dextrose (YEPD) medium at restrictive temperature (37°C). Aliquots were then taken at regular intervals over 5 hours for microarray analysis of transcript levels (Figure 1).

As hypothesized, the deletions of *WHI5* and *STB1* in the *cdc28-4* mutant substantially increased the mean transcript levels of the G1/S genes activated by SBF and MBF (Figures 1B, S1A, and S1B; p<2.2e-16 by paired t-test). However, most SBF/MBF targets were transcribed at high levels in the *cdk1 whi5Δ stb1Δ* mutant throughout the time course (Figure 1B and Figure 1C) and did not exhibit the pulsatile dynamics observed in wild-type cells. This observation was unexpected as the transcriptional repressors *NRM1/YHP1/YOX1* that were thought to mediate negative feedback loops also exhibited elevated transcript levels (Figure 1C). Moreover, the high-amplitude G1/S transcription triggered by SBF/MBF did not appear to pass through the TF network in the *cdk1 whi5Δ stb1Δ* mutant efficiently (Figure 1B). Despite the fact that the transcript levels of *HCM1* and *NDD1* were elevated as compared to the *cdc28-4* single mutant (Figure 1C), we did not observe corresponding increase in the expression levels of the majority of S/G2/M genes activated by Hcm1, SFF (Ndd1/Fkh2/Mcm1 complex), and Swi5/Ace2 (Figures 1B and S1A). Thus, in addition to inhibiting Whi5 and Stb1 to activate the G1/S transcriptional activating complexes SBF and MBF, Cln-CDKs appear to regulate other components of the TF network either directly or indirectly. These regulations by Cln-CDKs presumably contribute to the pulsatile dynamics of G1/S transcription and the serial activation of S/G2/M transcription that have been observed in the mutant cells lacking S-phase and mitotic cyclins (Figure S1C) (Orlando et al., 2008).

To facilitate comparison with further experiments described below, we repeated the experiments of *cdk1 whi5Δ stb1Δ* by synchronization with α-factor and obtained similar results.

### Anaphase-promoting complex (APC) prevents the accumulation of S/G2/M TFs in G1

We hypothesized that Cln-CDKs might promote either the activity or protein stability of downstream TFs activated by SBF/MBF. Indeed, it has been shown that the activity of Hcm1 is regulated by CDK phosphorylation (Landry et al., 2014). On the other hand, Nrm1, Yhp1, and Ndd1 appear to be substrates of APC^Cdh1^ (Edenberg et al., 2015; Ostapenko and Solomon, 2011; Sajman et al., 2015), which is an E3 ubiquitin ligase complex normally inactivated at G1/S transition by CDK phosphorylation (Huang et al., 2001; Jaspersen et al., 1999; Yeong et al., 2001; Zachariae et al., 1998). If Cdh1 is normally inactivated by CDK at the G1/S border, then the *cdc28-4* mutant cells should have constitutively active APC^Cdh1^, and thus APC^Cdh1^ substrates might not accumulate at the protein level.

We first examined the protein levels of these TFs in the *cdk1 whi5Δ stb1Δ* mutant. Cells carrying endogenously myc-epitope-tagged *NRM1*, *YHP1*, and *NDD1* were synchronized in G1 by α-factor at 25°C and then released at 37°C. The protein levels of Nrm1, Yhp1, and Ndd1 (collectively denoted as S/G2/M TFs below) were then measured by Western blot. In wild-type cells, these S/G2/M TFs were not detectable in early G1 and accumulated upon cell-cycle entry (Figure 2A). However, in the *cdk1 whi5Δ stb1Δ* mutant, these TFs only slowly accumulated and did not reach wild-type levels (Figures 2A and 2B), even though their transcript levels were comparable to wild-type levels (Figure 2B).

**Figure 2.**
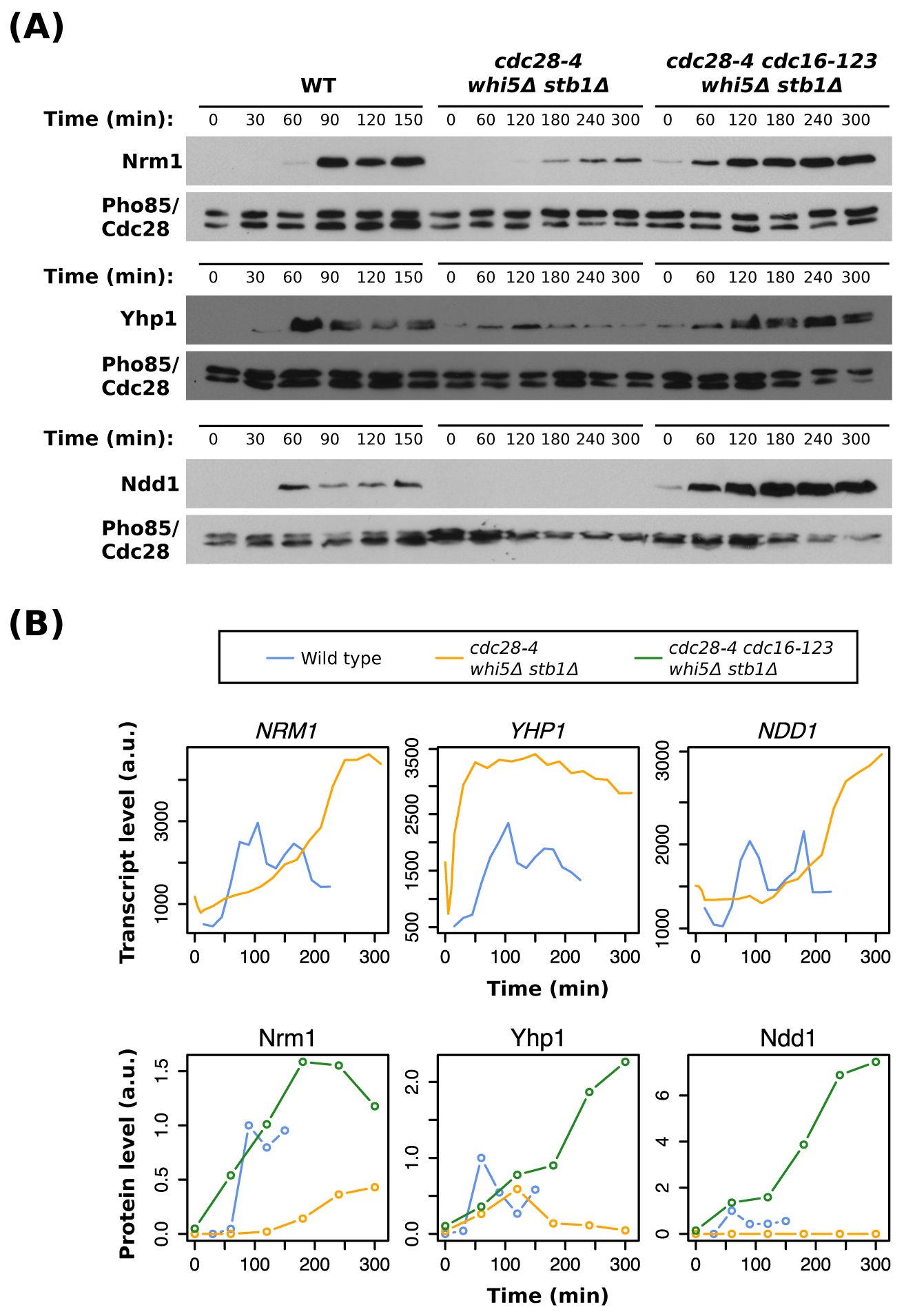
Inactivation of APC allows for the accumulation of S/G2/M TFs in the *cdc28-4* mutant. (A) Time-series Western blots of endogenously 13myc-tagged Nrm1, Yhp1, and Ndd1 in synchronized cell populations. Cells were synchronized by α-factor and released into YEP-dextrose (YEPD) medium at 37°C. Pho85 and Cdc28 detected by the α-PSTAIR antibody were used as loading control for quantitation. Representative results of three independent replicates are shown. (B) Line graphs showing transcript levels and protein levels of *NRM1, YHP1*, and *NDD1* in wild type (blue) (Simmons Kovacs et al., 2012), the *cdc28-4 whi5Δ stb1Δ* mutant (yellow), and the *cdc28-4 cdc16-123 whi5Δ stb1Δ* mutant (green). Cells were synchronized in early G1 and released into YEPD medium at 37°C. Transcript levels in cells released from elutriation were measured by microarray. Protein levels in cells released from α-factor and detected by Western blots shown in (A) were quantified and normalized to the Cdc28/Pho85 levels.

To investigate whether constitutively active APC^Cdh1^ in the *cdk1 whi5Δ stb1Δ* mutant prevents the accumulation of S/G2/M TFs, we wanted to introduce a *cdh1Δ* mutation into the *cdk1 whi5Δ stb1Δ* background. However, the *cdh1Δ whi5Δ* double mutations are synthetically lethal (Jorgensen et al., 2002), so we used a temperature-sensitive allele of *CDC16*, which encodes a component of APC, to perturb all APC activity (Irniger and Nasmyth, 1997). In the *cdk1 cdc16-123 whi5Δ stb1Δ* mutant, the accumulation of S/G2/M TFs was indeed restored after release at restrictive temperature compared to the *cdk1 whi5Δ stb1Δ* mutant (Figures 2A and 2B). These data suggest a model in which APC^Cdh1^ destabilizes S/G2/M TFs in G1 until its inactivation by G1/S cyclin-CDKs (Figure 3A).

**Figure 3.**
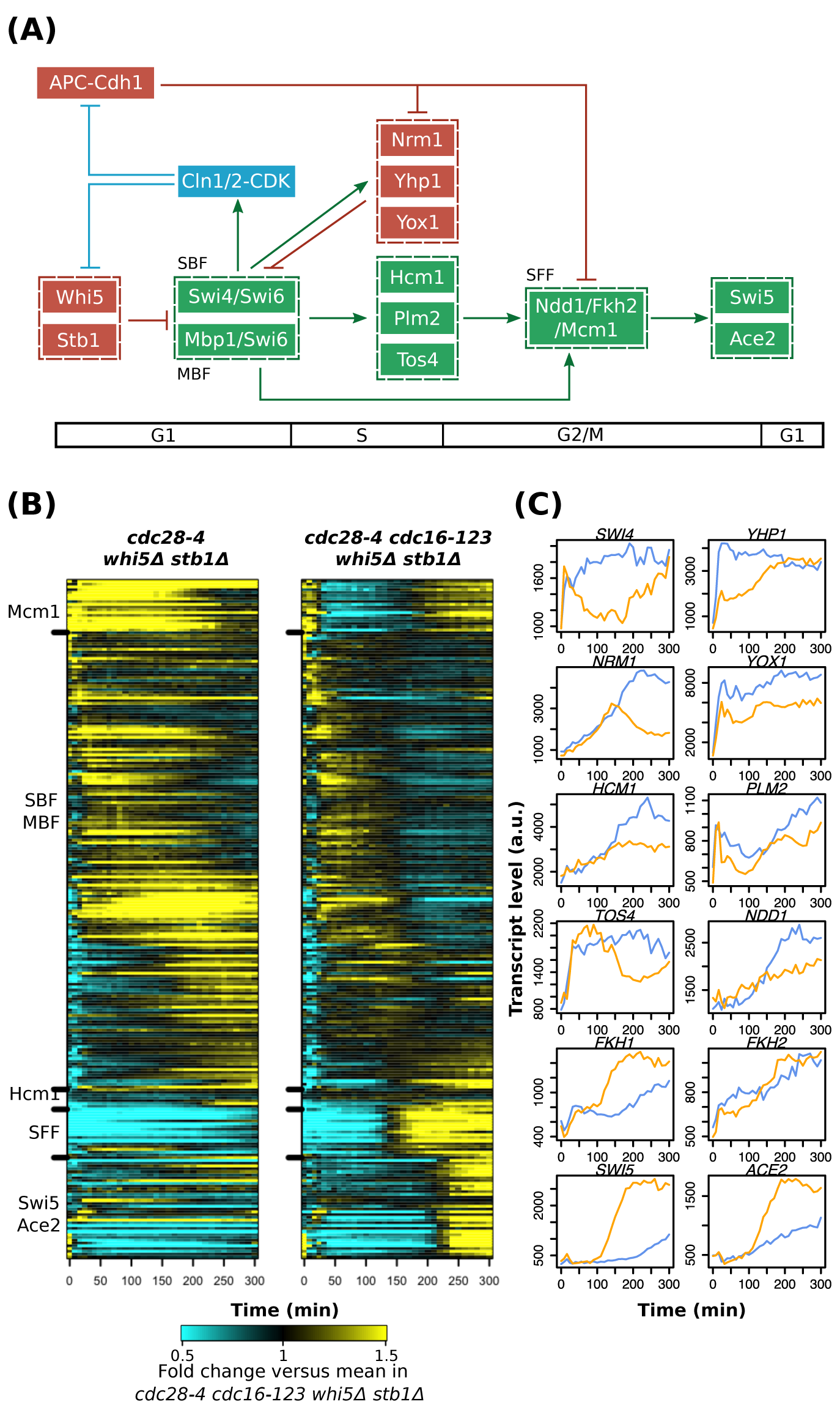
The inactivation of APC restores the transcriptional pulse and facilitates the transmission of the pulse through the network in *cdc28-4* mutant. (A) A revised network model indicating interactions between APC^Cdh1^, G1 cyclin-CDKs (blue) and relevant network TFs. Transcriptional activators and repressors are shown in green and red, respectively. Nodes are ordered horizontally by their approximate time of activation during the cell cycle. (B) Heat maps depicting transcript dynamics of the canonical genes regulated by the TF network (Table S1). Early G1 cells synchronized by α-factor were released into YEP-dextrose medium at 37°C. Transcript levels were measured by microarray and are depicted as fold change relative to mean in *cdk1 apc whi5Δ stb1Δ*. (C) Line graphs showing absolute transcript levels of the TF network components in the *cdk1 whi5Δ stb1Δ* (blue) and the *cdk1 apc whi5Δ stb1Δ* mutant (yellow) released from α-factor arrest. See also Figure S2.

### The inactivation of APC restores the generation and transmission of a transcriptional pulse by the TF network

The above results reveal that the dynamics of the TF network are inhibited at multiple levels by Whi5/Stb1/APC^Cdh1^. If Cln-CDKs promote robust cell-cycle transcription predominantly by mediating feedback to relieve these inhibitions, it should be possible to genetically restore the cell-cycle transcriptional program in the *cdk1 whi5Δ stb1Δ* mutant by further inactivating APC (Figure 3A). To test this hypothesis, we examined global transcript dynamics in the *cdc28-4 cdc16-123 whi5Δ stb1Δ* (denoted as *cdk1 apc whi5Δ stb1Δ* below) quadruple mutant by microarray. Early G1 cells were obtained by α-factor arrest at permissive temperature (25°C) and then released at restrictive temperature (37°C). Aliquots were then taken at regular intervals for 5 hours and subjected to microarray analysis (Figure 3).

In support of the hypothesis, the inactivation of APC activity restored much of the dynamics of cell-cycle transcription (Figures 3B and 3C). For a significant proportion of G1/S targets driven by SBF/MBF, a narrower transcriptional pulse was observed in the *cdk1 apc whi5Δ stb1Δ* mutant compared to the *cdk1 APC whi5Δ stb1Δ* mutant (Figures 3B, 3C, and S2A), which is consistent with the stabilization of repressors Nrm1 and Yhp1 (Figure 2). These results also support the notion that these transcriptional repressors are essential for producing pulsatile dynamics via negative feedback loops (Figure 3A). The lack of complete repression observed in a subset of SBF/MBF targets (Figure 3B; Figure S2A, see *CLN2* and *PCL1*) is consistent with the lack of Clb2-CDK activity, which has been established as an additional repressor for canonical SBF targets (Amon et al., 1993; Koch et al., 1996).

For S-phase targets driven by Hcm1, their coherent activation was still not observed in the *cdk1 apc whi5Δ stb1Δ* mutant (Figures 3B, 3C, and S2B). For example, the canonical Hcm1 targets *DSN1* and *CIN8* remained transcriptionally repressed in the *cdk1 apc whi5Δ stb1Δ* mutant (Figure S2B), further supporting previous findings that Hcm1 is post-transcriptionally regulated by CDK phosphorylations (Landry et al., 2014; Pramila et al., 2006). On the other hand, both *FKH1* and *FKH2* still exhibited weak oscillations in their transcript levels (Figure 3C), suggesting additional activating input other than Hcm1. Overall, the above results are reminiscent of the observations in the *hcm1Δ* mutant cells (Pramila et al., 2006).

While the dynamics of S-phase transcription remained partially impaired in the *cdk1 apc whi5Δ stb1Δ* mutant, G2/M transcription driven by SFF was coherently up-regulated (Figure 3B; Figure 3C, see *SWI5* and *ACE2*), likely due to the stabilization of the SFF component Ndd1 protein (Figure 2). Accordingly, we observed the subsequent activation of M/G1 transcription driven by Swi5/Ace2 upon APC inactivation (Figures 3B). Taken together, these observations support the idea that Cln-CDKs contribute to the robust transmission of the transcriptional pulse through the network by indirectly stabilizing Ndd1 (Figure 3A).

Although we did observe a transcriptional pulse moving through the network in the *cdk1 apc whi5Δ stb1Δ* mutant cells (Figures 3B and 3C), we did not observe a robust second pulse in most of the program except the early-G1 transcription activated by Mcm1 (Figure 3B). Because of the stabilization of transcriptional repressors Nrm1 and Yhp1 in cells carrying the APC mutant allele *cdc16-123* (Figure 2), the inhibition of a second cycle of SBF/MBF-mediated transcription was expected.

### The inhibition of Whi5/Stb1/APC in the *cdc28-4* mutant cells restores phase-specific transcription at high amplitude

We next wanted to determine the extent to which the cell-cycle transcriptional program was restored by the simultaneous inhibition of Whi5/Stb1/APC in the *cdc28-4* background. First, we asked whether the transcript levels driven by the TF network were comparable between the *cdk1 apc whi5Δ stb1Δ* mutant and wild-type cells. As shown in Figure 4A, the *cdk1 apc whi5Δ stb1Δ* mutant cells were able to activate G1/S transcription (SBF/MBF targets) and G2/M transcription (SFF targets) at levels similar to wild-type cycling cells at 37°C, while partial restoration of the S-phase transcription (Hcm1 targets) and M/G1 transcription (Swi5/Ace2 targets) were also observed.

**Figure 4.**
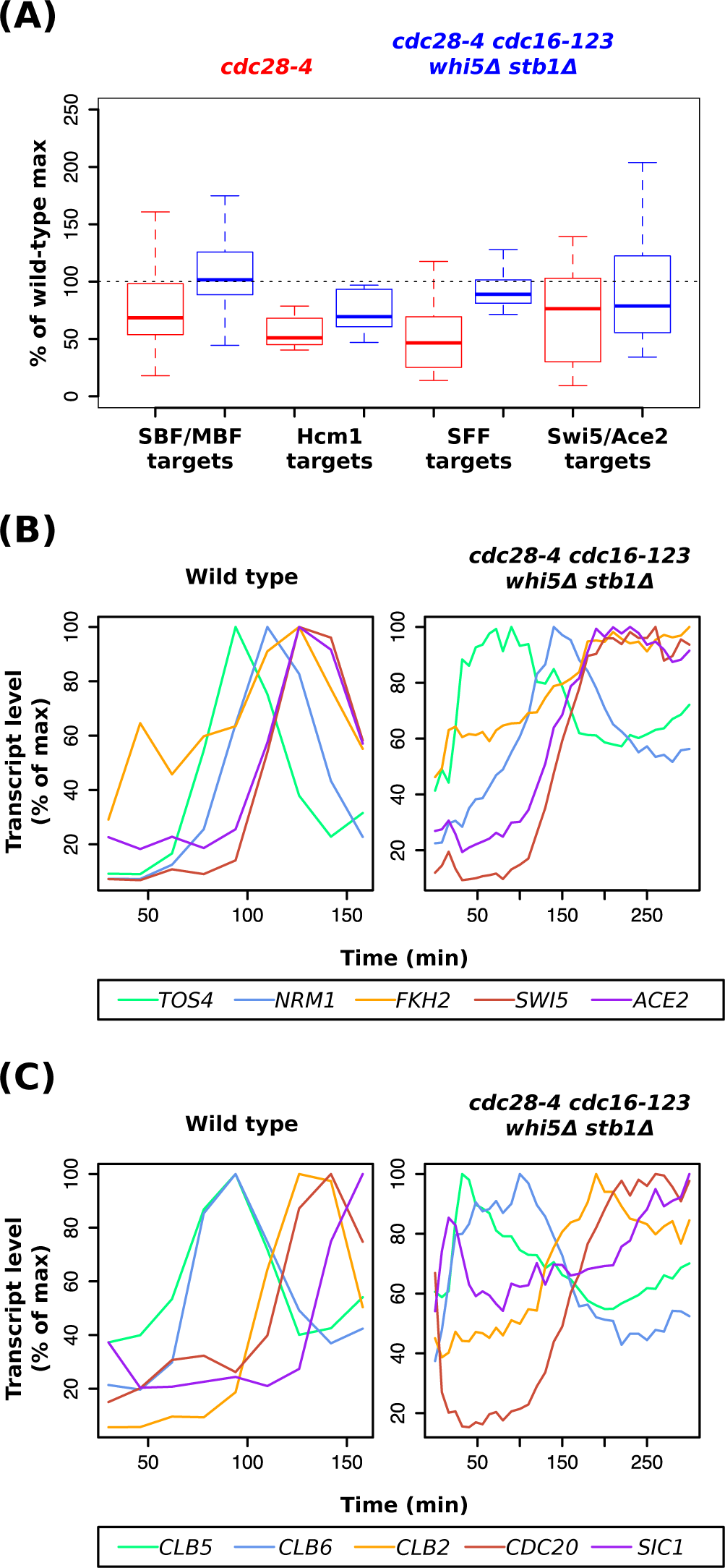
The *cdc28-4 cdc16-123 whi5Δ stbΔ* cells trigger a temporally ordered transcriptional program at high amplitude. (A) Box plots depicting maximal expression levels of the TF network targets in the *cdc28-4* experiments (Simmons Kovacs et al., 2012) and the *cdc28-4 cdc16-123 whi5Δ stb1Δ* experiments. The average from two independent replicates is plotted as percent of wild-type control at 37°C (Simmons Kovacs et al., 2012). Whiskers extend to 1.5 times interquartile range from the box. Outliers in the data are not shown. (B)(C) Line graphs showing transcript levels of network TFs (B) or CDK regulators (C) in experiments shown in (A). Transcript levels are plotted as percentage of maximal level for each gene in individual time courses. See also Figures S3 and S4.

Next, we asked whether the ordering of the serial activation of network TFs was still conserved. To this end, we directly compared the transcript dynamics of those TFs that exhibited oscillatory behaviors in the *cdk1 apc whi5Δ stb1Δ* mutant cells (Figure 3C) to their dynamics in wild-type cells. In support of a model in which the TF network can transmit a transcriptional pulse, a subset of TF network components were activated in the same order in both wild type and the *cdk1 apc whi5Δ stb1Δ* mutant cells with similar peak-to-trough ratios (Figure 4B). Strikingly, the phase-specific transcription of CDK regulators, including that of the S-phase cyclins *CLB5/6*, the mitotic cyclin *CLB2*, the APC coactivator *CDC20*, and the B-cyclin-CDK inhibitor *SIC1*, was also conserved in the *cdk1 apc whi5Δ stb1Δ* mutant (Figure 4C).

These findings support the idea that the TF network in concert with Cln-CDKs contributes to cell-cycle progression by generating a high-amplitude, properly ordered cell-cycle transcriptional program.

To expand our analyses to the dynamics of global cell-cycle transcription, we utilized a high-confidence periodic gene set (Bristow et al., 2014) for further analysis. We excluded genes in the environmental stress response to avoid the transcript dynamics induced by the temperature shift during the experiments (Gasch et al., 2000). We found that the inhibition of Whi5/Stb1/APC in the *cdc28-4* background greatly improved both the amplitude and phase-specific ordering of this periodic transcriptional program containing 857 genes (Figure S3 and Table S2).

Taken together, the above data support a model in which the inhibition of Whi5/Stb1/APC^Cdh1^ by G1 cyclin-CDKs is sufficient to allow the TF network to trigger a large program of cell-cycle transcription at high amplitude. However, the speed at which the pulse was propagated was still slower in the *cdk1 apc whi5Δ stb1Δ* mutant cells compared to wild type (Figures 4B, 4C, and S3), suggesting additional mechanisms by which CDKs promote the dynamics of the global cell-cycle transcription.

### A large program of cell-cycle transcription continues in cells overexpressing hyperstable B-cyclin-CDK inhibitor, Sic1

Interestingly, we noticed that the deletion of *WHI5* or *STB1* in the *cdc28-4* background triggered bud emergence in early G1 cells even at restrictive temperature (Figure S1D). Further deletions of *PCL1/PCL2* (G1 cyclins for the CDK Pho85) severely delayed or inhibited budding (data not shown), suggesting that the bud emergence in these mutants were dependent on the Pcl1/2-Pho85 kinase activity (Moffat and Andrews, 2003). Double deletions of *WHI5* and *STB1* resulted in the earliest bud emergence after release (Figure S1E), suggesting the strongest derepression of *PCL1/PCL2* among the SBF/MBF targets. Finally, the *cdc28-4/cdk1 apc whi5Δ stb1Δ* quadruple mutant triggered re-budding cycles of elongated buds, suggesting the lack of mitotic Clb2-CDK activity that inhibits bud polarity (Figure S4D).

However, we did observe that a fraction of cells underwent DNA replication and spindle pole body duplication in the *cdc28-4/cdk1 apc whi5Δ stb1Δ* mutant after several hours at restrictive temperature (Figures S4A and S4C). The DNA replication was blocked by the overexpression of hyperstable Sic1 in these cells (Figure S4B) (Verma et al., 1997), suggesting that the protein product encoded by the *cdc28-4* allele can still be weakly activated by S-phase and/or M-phase B-cyclins (Clbs) at restrictive temperature. Given the surprising finding that enough Clb-CDK activity remains in some *cdc28-4 apc whi5Δ stb1Δ* mutant cells to drive DNA replication, and the suggestion that small amounts of residual Clb could drive the transcriptional program in the *clbΔ* mutant cells (Rahi et al., 2016), we wanted to validate the above findings and further test our network model (Figure 3A) in additional mutants where Clb-CDKs are more fully inhibited.

The inhibition of both S-phase and mitotic Clb-CDK activity by Sic1 has been confirmed by genetic and protein-protein interaction (Breitkreutz et al., 2010; Cross et al., 2007; Schreiber et al., 2012). The non-phosphorylatable SicΔ3P protein is hyperstable and delays cell-cycle progression when expressed at physiological level (Cross et al., 2007), while its overexpression blocks DNA replication and arrests the cell cycle (Verma et al., 1997). In current quantitative models of the budding yeast cell cycle, the overexpression of hyperstable SicΔ3P eliminates all Clb-CDK activity (Chen et al., 2004; Kraikivski et al., 2015). In summary, the *GAL-SICΔ3P* cells are presumably arrested without residual Clb-CDK activity, while Cln1/2-CDK activity remains constitutively high (Schwob et al., 1994; Verma et al., 1997).

To assay the global transcript dynamics, early G1 cells carrying the *GAL-SICΔ3P* construct were collected by centrifugal elutriation and then released into YEP-galactose (YEPG) media to induce overexpression. Samples were taken every 10 minutes for time-series microarray (Figure 5). The physical cell-cycle arrest was confirmed by monitoring re-budding cycles, which phenocopy the *clb1-6Δ* mutant (data not shown) (Haase and Reed, 1999). Consistent with previous findings (Orlando et al., 2008), a large program of cell-cycle transcription continued in the *GAL-SICΔ3P* cells lacking Clb-CDK activity (Figure 5A). Notably, the generation and transmission of a transcriptional pulse were observed for the majority of the cell-cycle genes (Figure 5A). These observations further support the idea that G1 cyclin-CDKs can not only activate G1/S transcription but also enhance the global dynamics of cell-cycle transcription. In support of the network model (Figure 3A), we also observed a transcriptional pulse propagated through the TF network in the *GAL-SICΔ3P* cells with temporal ordering identical to wild type (Figure 5B).

**Figure 5.**
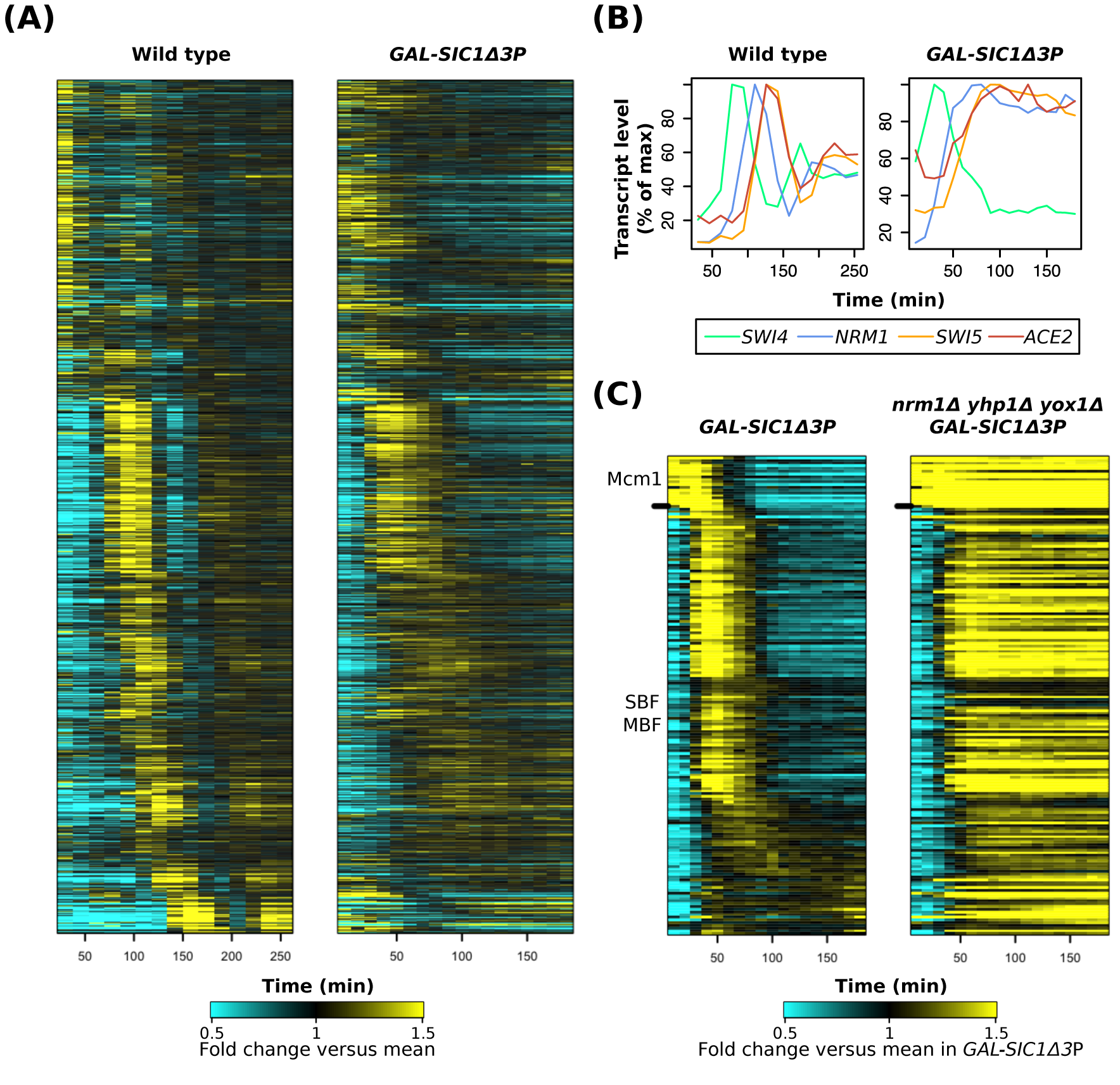
A well-ordered transcriptional program is maintained in cells overexpressing hyperstable B-cyclin-CDK inhibitor Sic1. (A) Heat maps showing transcript dynamics of 857 cell-cycle genes (Table S2) in wild type (Orlando et al., 2008) and the *GAL-SIC1Δ3P* strain. Early G1 cells obtained by elutriation were released into YEP-galactose medium at 30°C for microarray analysis. Transcript levels are expressed as fold change relative to mean in individual datasets. (B) Line graphs showing transcript levels of network TFs in wild type and the *GAL-SIC1Δ3P* cells. Transcript levels are expressed as percentage of maximal level in individual time courses. (C) Heat maps showing transcript dynamics of Mcm1 and SBF/MBF targets (Table S1) in the *GAL-SIC1Δ3P* and the *GAL-SICΔ3P nrm1Δ yhp1Δ yox1* strains. Cells were synchronized in G1 by elutriation and released into YEP-galactose medium at 30°C. Transcript levels are expressed as fold change relative to mean in the control *GAL-SICΔ3P* experiment. See also Figure S5.

To confirm that the G1/S transcriptional pulse in the *GAL-SICΔ3P* cells is generated by negative feedback loops from transcriptional repressors in the TF network rather than residual Clb2 activity (Figure 3A), we further deleted *NRM1, YHP1*, and *YOX1* and assayed transcript dynamics. In the *GAL-SICΔ3P* cells, a majority of Mcm1 targets and SBF/MBF targets were robustly attenuated (Figures 5C and S5). In the *GAL-SICΔ3P nrm1Δ yhp1Δ yox1Δ* cells, all Mcm1 targets and SBF/MBF targets were activated and then remained highly expressed throughout the time course (Figures 5C and S5). The loss of pulsatile dynamics supports the critical role of these transcriptional negative feedback loops in generating transcriptional pulses in a variety of conditions, including the *clb1-6Δ* and the *cdk1 apc whi5Δ stb1Δ* mutant cells. Similarly to the above results in the *cdk1 apc whi5Δ stb1Δ* mutant (Figure 3B), we did not observe a robust second pulse for SBF/MBF targets in the *GAL-SICΔ3P* cells (Figure 5C), suggesting the stabilization of transcriptional repressors Nrm1/Yhp1/Yox1 via Cln-CDK-dependent APC inactivation.

Taken together, these results from the *GAL-SICΔ3P* experiments further support a model in which G1 cyclin-CDKs and the TF network function in an integrated network to generate a high-amplitude cell-cycle transcriptional program after cell-cycle commitment (Figure 3A).

## DISCUSSION

Although the oscillations of CDK and APC activity have been thought as the central oscillator that dictates phases of other cell-cycle oscillations, there is increasing evidence in recent years that global cell-cycle transcription can continue semi-autonomously without periodic CDK-APC activity. Collectively, previous studies pointed to a model in which both CDKs and a transcription factor (TF) network have critical roles in controlling global cell-cycle transcription, but that oscillating input from CDK is not required to generate transcriptional dynamics (Bristow et al., 2014; Cho et al., 2017). The most compelling evidence for this model comes from the observations that temporally ordered, high-amplitude transcript dynamics could still be observed in budding yeast cells arrested with constitutive levels of CDK activity. In cells lacking all CDK activity, only low-amplitude transcriptional oscillations were observed in a fraction of the program, and they displayed an increased period length when compared to wildtype cells (Simmons Kovacs et al., 2012). These findings suggest a role of CDKs in promoting transcriptional oscillations, yet the mechanism was not understood.

Here we propose a model in which G1 cyclin-CDKs contribute to the generation of the cell-cycle transcriptional program by the TF network via multiple molecular mechanisms. First, G1 cyclin-CDKs inactivate transcriptional corepressors Whi5/Stb1 to trigger the high-amplitude transcriptional activation of SBF/MBF targets (Figure 1). Second, the partial inactivation of APC^Cdh1^ by G1 cyclin-CDKs stabilizes network transcriptional repressors (Nrm1 and Yhp1) and coactivator Ndd1, which are crucial for robust oscillations of the TF network. Particularly, the transcriptional repressors Nrm1, Yhp1, and Yox1 mediate negative feedback to truncate the G1/S transcriptional activation into a transcriptional “pulse” (Figure 3 and Figure 5). A chain of transcriptional activators downstream of SBF/MBF then transmits the pulse and serially activates S/G2/M transcription.

Consistent with the above model and previous findings, we provide further evidence that S-phase and mitotic Clb-CDK activities are largely dispensable for generating and transmitting the transcriptional pulse that drives cell-cycle transcription. In cells overexpressing hyperstable Sic1 that specifically inhibit Clb-CDK activities, a robust G1/S transcriptional pulse can still be generated, and that the serial activation of S/G2/M transcription persist despite the physical G1/S arrest (Figure 5). We also demonstrate that the transcriptional repressors

Nrm1/Yhp1/Yox1 are necessary for the attenuation of early-cell-cycle transcription in the absence of Clb-CDK activities, supporting the roles of these TFs in generating pulsatile transcript dynamics in either wild-type or cyclin mutant cells.

We have demonstrated previously that oscillations of the cell-cycle transcriptional program can be uncoupled from CDK oscillations and from cell-cycle progression, and that S-phase and M-phase checkpoints can halt the dynamics of cell-cycle transcription when cellcycle progression is perturbed (Bristow et al., 2014; Haase and Reed, 1999; Orlando et al., 2008; Simmons Kovacs et al., 2012). Our findings here suggest that G1 is also an important phase for coordinating the oscillations of CDK activity and the cell-cycle transcriptional program (Figure 6A). Specifically, transcriptional corepressors Whi5/Stb1 directly inhibit the G1/S transcriptional activators SBF/MBF, thus indirectly inhibiting the transcription of S/G2/M TFs activated by SBF/MBF. Furthermore, APC^Cdh1^ destabilizes several S/G2/M TFs as well as S-phase and mitotic cyclins (Clbs). Finally, the stoichiometric inhibitor Sic1 binds to and inhibits all Clb-CDK activities. Thus, the combined activities of Whi5/Stb1, APC^Cdh1^, and Sic1 inhibit both oscillations of CDK activity and the TF network at transcriptional and post-translational levels in G1 phase (Figure 6A). This idea is similar to the “G1 attractor” in a previously proposed model of the budding yeast cell-cycle network (Li et al., 2004).

**Figure 6.**
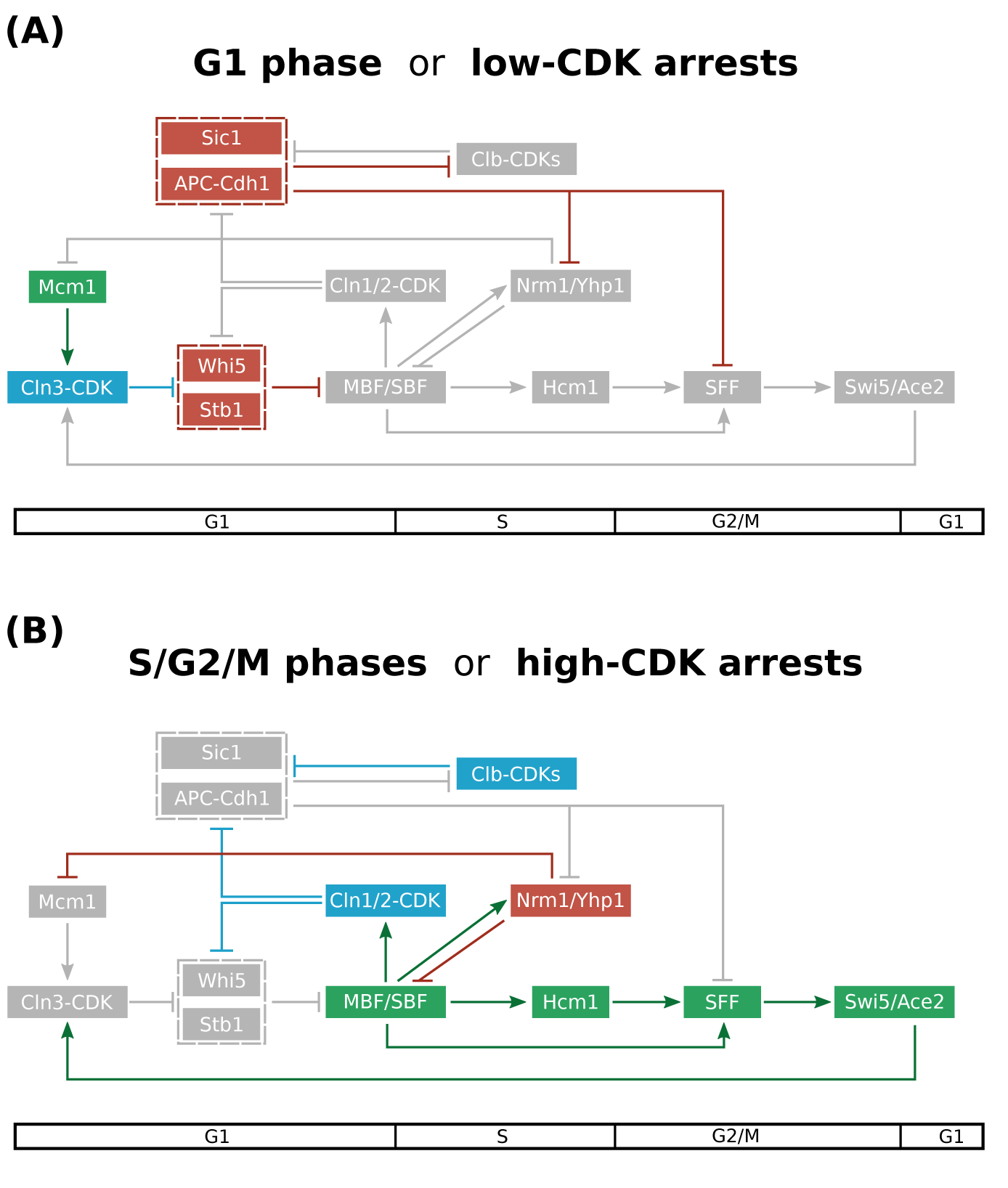
An integrated network model for the global control of cell-cycle transcription. Nodes are ordered horizontally by their approximate time of activation during the cell cycle. (A) In early G1, both CDKs (blue) and the TF network (green) are globally inhibited by G1 inhibitors (red), including Whi5/Stb1, APC^Cdh1^, and Sic1. (B) In S/G2/M phases, the CDK-dependent inhibition of Whi5/Stb1/APC^Cdh1^/Sic1 allows the robust oscillations of the TF network.

In wild-type cells, these inhibitions are likely maintained until increasing cell size dilutes Whi5, which results in the initial expression of *CLN1/2* in a “feedback-first” mechanism (Eser et al., 2011; Schmoller et al., 2015). Subsequently, Cln1/2-CDKs mediate feedback loops to inactivate these G1 inhibitors (Whi5/Stb1/APC^Cdh1^/Sic1) and trigger entry into S/G2/M phases (Figure 6B). This release of inhibition allows the TF network to generate a G1/S transcriptional pulse and transmit the pulse to generate global cell-cycle transcription, including the temporally ordered transcription of Clbs. Once transcriptionally activated by the TF network, Clb-CDKs trigger S-phase and mitotic events, while also mediating feedback to modulate the amplitude and timing of cell-cycle transcription by inhibiting SBF-mediated transcription and activating SFF-mediated transcription (Amon et al., 1993; Koch et al., 1996; Pic-Taylor et al., 2004; Reynolds et al., 2003). Furthermore, while the inactivation of APC^Cdh1^ and Sic1 can be initiated by Cln-CDKs during G1/S transition, Clb-CDKs later reinforce and maintain the inhibition of APC^Cdh1^ and Sic1 in S/G2/M phases (Figure 6B). Finally, Cdc14 phosphatase released by mitotic exit pathways coordinates the destruction of Nrm1, Yhp1, Yox1, and Clb2 with the reactivation of Whi5/Stb1 (via dephosphorylation) to maintain the repression of SBF/MBF-regulated genes in the next G1 phase (Taberner et al., 2009; Wagner et al., 2009). These layers of repression ensure that high-amplitude transcription is activated once-and-only-once per cycle in wild-type cells. Interestingly, the transcriptional pulse could eventually be transmitted back to regulate the expression of G1 cyclin *CLN3* through network TFs Swi5/Ace2, providing an additional mechanism for promoting a new pulse in the next cycle (Di Talia et al., 2009).

Significantly, this model provides a unifying explanation for the transcriptomic dynamics in a broad range of budding yeast mutant cells arrested in the cell cycle (Figure 6). In cells arrested with low CDK activity, such as the *cdc28-4* mutant cells (Simmons Kovacs et al., 2012) or the *cln clb* mutant cells (Rahi et al., 2016), the activity of Whi5/Stb1/APC^Cdh1^ then prevents the initiation of a robust transcriptional pulse (Figure 6A). In cells arrested with constitutive G1 Cln-CDK or mitotic Clb-CDK activity, such as the *clb1-6Δ, cdc20Δ*, and *cdc14-3* mutants (Bristow et al., 2014; Cho et al., 2017; Orlando et al., 2008; Rahi et al., 2016), the TF network can continue to trigger a subset of the cell-cycle transcriptional program due to the CDK-dependent inhibition of Whi5/Stb1/APC^Cdh1^ activity (Figure 6B). Finally, we demonstrate in this study that genetically perturbing G1 inhibitors or overexpressing Clb-CDK inhibitor can also uncouple the dynamics of the TF network from the oscillation of CDK activity (Figures 3 and 5).

Given the topological conservation of at least part of the cell-cycle networks (Cross et al., 2011; Johnson and Skotheim, 2013), findings for the budding yeast cell cycle will likely provide further insight into the cell-cycle regulatory mechanisms in mammalian cells. Here we report that the inactivation of APC^Cdh1^ by CDK phosphorylations is necessary for the attenuation of G1/S transcription and the activation of G2/M transcription. Interestingly, the G1/S transcriptional repressors E2F7/E2F8 and the mitotic transcriptional activator FoxM1 in mammalian cells are also APC^Cdh1^ substrates (Cohen et al., 2013; Park et al., 2008). Furthermore, it has been proposed that the irreversible inactivation of APC/C^Cdh1^ is the commitment point for mammalian cell cycle (Cappell et al., 2016). Similar genetic-genomic studies will be needed in order to dissect the contributions of CDKs, APC/C, checkpoint kinases, and a transcriptional network to the dynamics of periodic cell-cycle transcription in higher eukaryotes. Finally, we expect the phase-specific, multi-layered inhibition of the transcription factor network to be a general mechanism that restricts genome-wide transcriptional programs, such as during mammalian cell cycle or circadian oscillations, to one pulse per cycle.

## EXPERIMENTAL PROCEDURES

Requests of reagents and further information may be directed to the corresponding author Steven B. Haase.

### Yeast strains and cell culture synchronization

All strains are derivatives of S. *cerevisiae* BF264-15D (*ade1 his2 leu2-3,112 trp1-1a)*. Additional genotypes can be found in Table S3. Gene deletions and epitope tagging were carried out by standard yeast methods (Longtine et al., 1998). Strain K4438 (W303 *cdc16-123*) was kindly provided by Kim Nasmyth (Irniger and Nasmyth, 1997) and outcrossed with BF264-15D for 5 times before crossing into SBY2356 (*cdc28-4 whi5Δ stb1Δ BAR1*) to obtain SBY2395 (*cdc28-4 cdc16-123 whi5Δ stb1Δ bar1*).

Yeast cultures were grown in standard YEP medium (1% yeast extract, 2% peptone, 0.012% adenine, 0.006% uracil supplemented with 2% sugar). For centrifugal elutriation of temperature-sensitive strains carrying *cdc28-4* and/or *cdc16-123* alleles, cultures were grown to mid-log phase in YEP-galactose (YEPG) medium at 30°C. Elutriated early G1 cells were then resuspended in YEP-dextrose (YEPD) medium at 37°C. For α-factor arrest of temperature-sensitive strains, cultures were grown in YEPG medium at 25°C and incubated with 50 ng/ml α-factor for 140 minutes. Synchronized cultures were then resuspended in YEPD medium at 37°C. For *GAL-SIC1Δ3P* strains, cultures were grown to mid-log phase in YEP-sucrose (YEPS) medium at 30°C. Elutriated early G1 cells were then resuspended in YEP-galactose (YEPG) medium at 30°C for time-course experiments. Aliquots were taken at each time point and subsequently assayed by microarray or Western blots.

### RNA extraction and microarray assay

Total RNA was isolated by standard acid phenol protocol (2001). Samples were submitted to Duke Center for Genomic and Computational Biology Microarray Facility for labeling, hybridization, and image collection. mRNA was amplified and labeled by Ambion MessageAmp Premier kit (Ambion Biosystems) and hybridized to Yeast Genome 2.0 Array (Affymetrix).

### Compilation of canonical targets of the TF network

We compiled a list of canonical genes regulated by the network TFs for analyses. This list of genes and their microarray probe IDs is provided in Table S1. As described below, we considered 5 major groups of co-regulated genes: Mcm1 targets, SBF/MBF targets, Hcm1 targets, SFF targets, and Swi5/Ace2 targets. Each group contained unique genes, where genes with multiple regulations were only assigned to one cluster.

The Mcm1 cluster repressed by Yhp1/Yox1 has been previously reported (Pramila et al., 2002). We further excluded genes that were not co-expressed with Mcm2-6 in the wild type datasets (Orlando et al., 2008). This resulted in 18 genes that are coherently expressed in early G1.

The SBF/MBF targets are expressed at the G1/S transition and have been previously reported by Ferrezuelo et al. (2010). *HO* was excluded because its transcript level peaked at M/G1 rather than G1/S transition in our wild type datasets (Orlando et al., 2008; Simmons Kovacs et al., 2012). We further restricted our analyses to the 161 genes that have uniquely mapped probes in the microarray.

We defined the Hcm1 targets by two criteria: (1) they have documented expression evidence on YEASTRACT (Teixeira et al., 2014); AND (2) their transcript dynamics in the wild type datasets (Orlando et al., 2008) are clustered together with *CIN8* by affinity propagation (Frey and Dueck, 2007).

We defined the SFF (Ndd1/Fkh2/Mcm1 complex) targets with similar criteria: (1) they have documented DNA binding OR expression evidence for Fkh2 AND Mcm1 on YEASTRACT; AND (2) their transcript dynamics in the wild type datasets (Orlando et al., 2008) are clustered together with *CLB2* by affinity propagation. The clustering identified two groups of genes with peak expression in M phase. We excluded the cluster containing *CDC20* that exhibited an early minor peak in the wild type datasets to avoid complex regulations by factors other than SFF. The remaining cluster contains 17 genes that partially overlapped with previously reported CLB2 cluster or SFF targets (Sajman et al., 2015; Spellman et al., 1998; Zhu et al., 2000).

The Swi5/Ace2 targets have been previously reported (Di Talia et al., 2009). We further excluded genes whose transcript dynamics in wild-type cells are not clustered together with *SIC1* by affinity propagation.

*CLB1-6, WHI5, STB1, NRM1*, and *YOX1* are excluded for analyses involving their deletion mutants. In the microarray analysis, one representative and uniquely mapped probe was used for each gene.

### Protein isolation and Western blotting

Cell pellets were washed with ice-cold water and resuspended in TCA extraction buffer (1.4 M sorbitol, 25 mM Tris-HCl pH7.5, 20 mM NaN3, 2 mM MgCl2, and 15% TCA). Cell lysis was achieved by vortexing with glass beads at 4°C for 10 minutes. Pellets were collected by centrifuge and resuspended in Thorner buffer (8 M Urea, 5% SDS, 40 mM Tris-HCl pH 6.8, 0.1 mM EDTA, 0.4 mg/ml Bromophenol Blue, and 1% β-ME). Samples were titrated with 1 M Tris, heated at 42°C for 5 minutes, separated by SDS-PAGE, and transferred to Immobilon-P PVDF membrane (Millipore) for antibody probing. Western blotting was performed using the following antibodies: mouse anti-PSTAIR (Abcam), mouse anti-c-Myc clone 9E10 (Santa Cruz Biotechnology), anti-mouse IgG-HRP (Cell Signaling), and anti-rabbit IgG-HRP (Cell Signaling).

### Microscopy

Cells were fixed in 2% paraformaldehyde for 5 minutes at room temperature, washed with PBS, and then resuspended in 30% glycerol for mounting on glass slides. All imaging was performed on Zeiss Axio Observer.

### Flow cytometry

Cells were prepared for flow cytometric analysis using SYTOX Green staining as described (Haase and Reed, 2001). Graphs were generated using the FlowViz package in Bioconductor in R.

### Normalization of microarray data

Previously published datasets used in this study are GEO: GSE8799, GEO: GSE32974, and GEO: GSE49650. All CEL files analyzed in this study were normalized together using the dChip method from the Affy package in Bioconductor as described previously (Bristow et al., 2014).

### Data availability

Newly generated array data and the normalized data have been submitted to GEO: GSE75694.

## AUTHOR CONTRIBUTIONS

Conceptualization, S.B.H., C.C., and C.M.K.; Methodology and Formal Analysis, C.C., C.M.K.; Writing - Original Draft, S.B.H. and C.C.; Writing - Review & Editing, C.M.K.

## ACKNOWLEDGEMENTS

We thank Daniel J. Lew and Adam R. Leman for helpful discussions and critical reading of the manuscript. We also thank Kim Nasmyth for kindly supplying the *cdc16-132* allele. This work was supported by the Defense Advanced Research Projects Agency grant #D12AP00025. There is no conflict of interest over the research findings shared in this manuscript.

---

**Figure S1.**
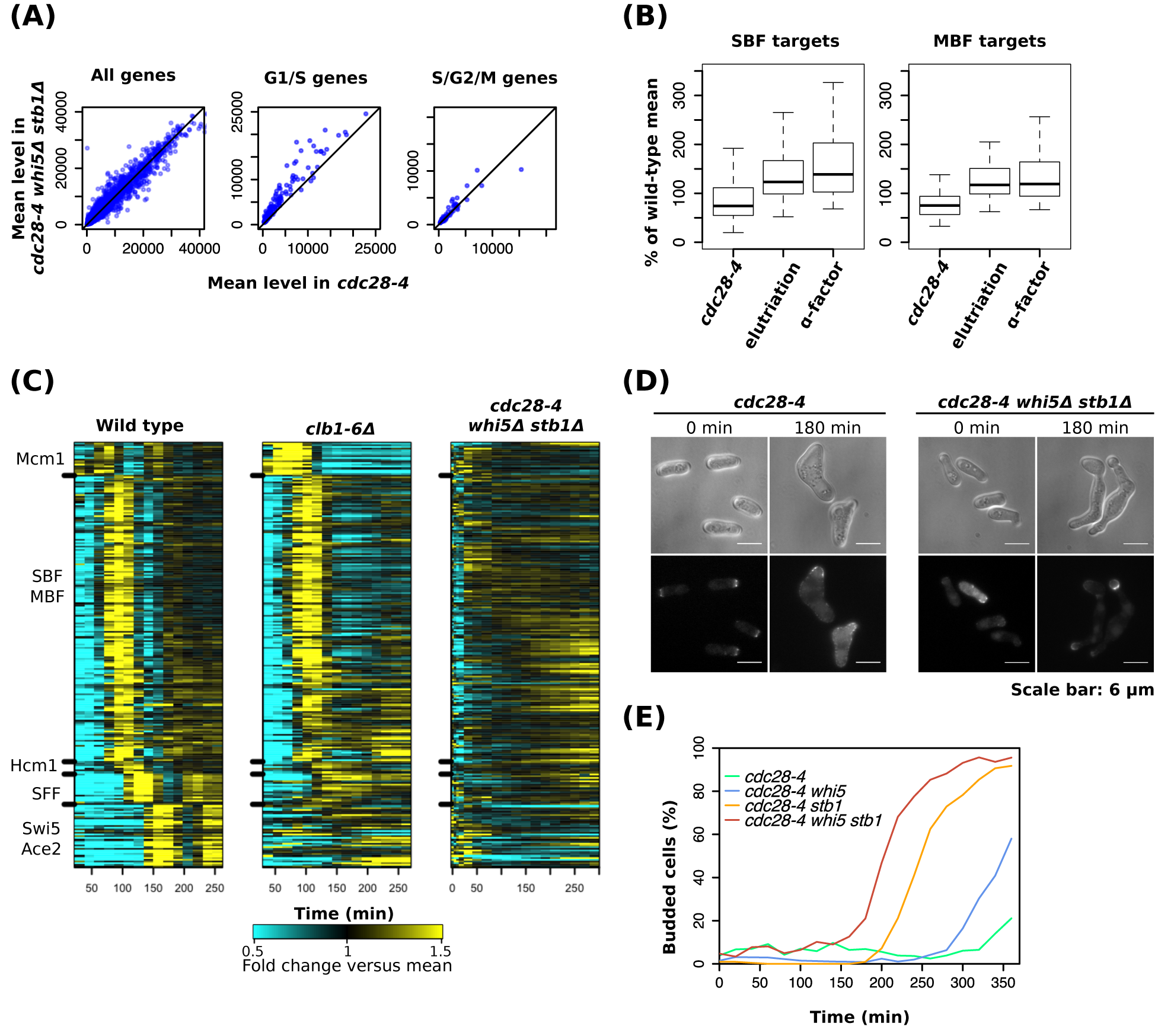
Additional analyses related to the *cdc28-4 whi5Δ stb1Δ* experiments (related to Figure 1). (A) Scatter plots showing mean transcript levels of all genes, G1/S genes (SBF/MBF targets), or S/G2/M genes (Hcm1/SFF/Swi5/Ace2 targets) during the time courses in indicated strains. Average of results from two independent replicates are shown. (B) Box plots depicting mean expression levels of the SBF/MBF targets in the *cdc28-4* experiment (Simmons Kovacs et al., 2012) and the *cdc28-4 whi5Δ stb1Δ* experiments synchronized by elutriation or α-factor. The groups of SBF and MBF targets were reported in the previous study (Ferrezuelo et al., 2010). The average from two independent replicates is plotted as percent of wild-type control at 37°C (Simmons Kovacs et al., 2012). Whiskers extend to 1.5 times interquartile range from the box. Outliers in the data are not shown. (C) Heat maps showing transcript dynamics of the canonical genes regulated by the TF network in indicated strains. Representative results of wild type, the *clb1-6Δ* experiment released from elutriation at 30°C (Orlando et al., 2008), or the *cdc28-4 whi5Δ stb1Δ* experiment released from elutriation are shown. Transcript levels are depicted as fold change versus mean in individual dataset. (D) Microscopic images of the *cdc28-4* and *cdc28-4 whi5Δ stb1Δ* mutants. Cells were synchronized in G1 by centrifugal elutriation, released into YEPD medium at 37°C, and fixed at indicated time points for subsequent imaging. The bud emergence in *cdc28-4 whi5Δ stb1Δ* is confirmed by the formation of septin rings (*CDC3-mCherry*) shown in the bottom panels. (E) The budding curves of various *cdc28-4* strains carrying *WHI5* and/or *STB1* deletion after released from elutriation into YEPD medium at 37°C. Cells with visible restrictions on the cell body were counted as budded cells. The *CDC3-mCherry* marker was not used for scoring.

**Figure S2.**
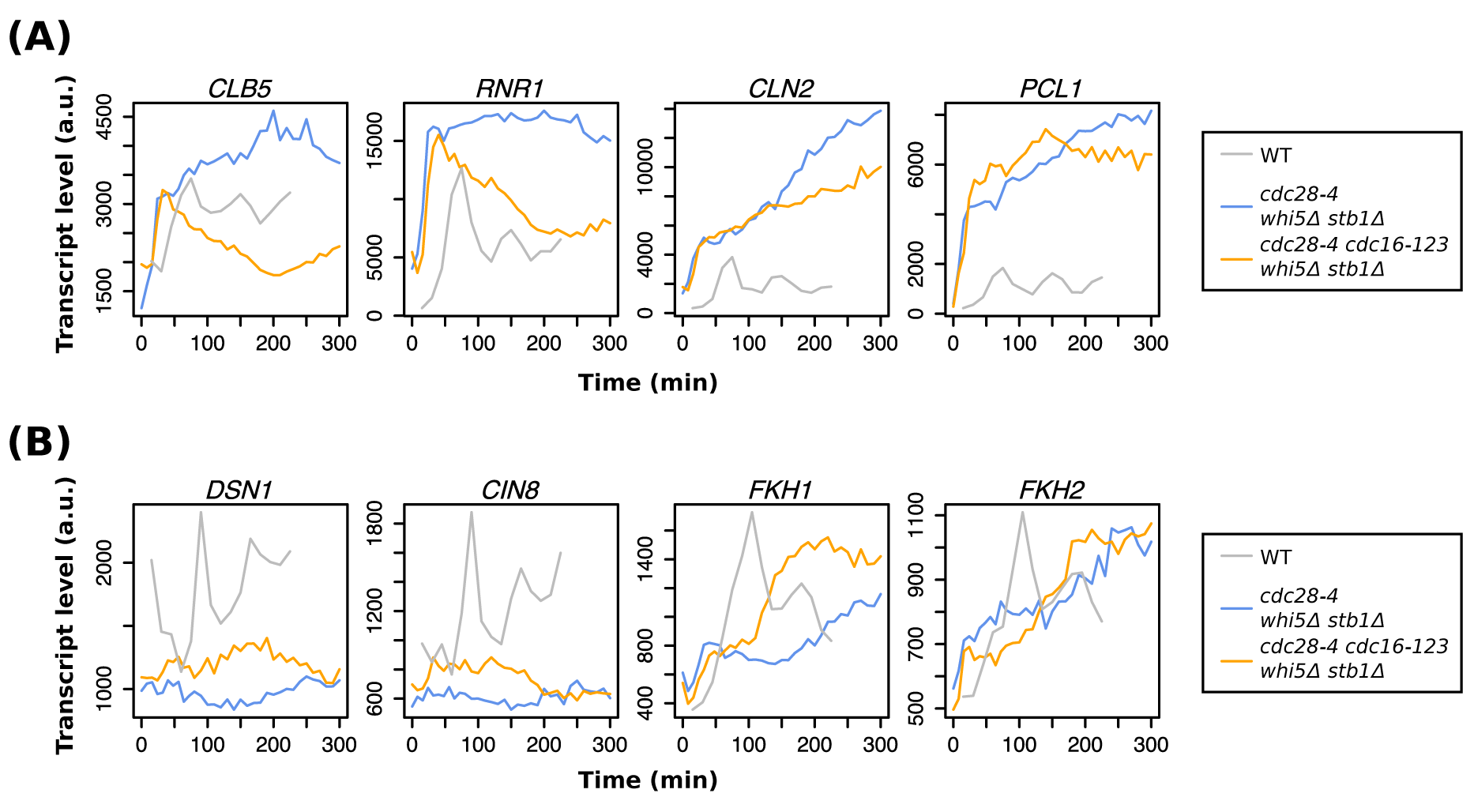
Additional analyses related to the *cdc28-4 whi5Δ stbΔ* and *cdc28-4 cdc16-123 whi5Δ stbΔ* experiments (related Figure 3). (A) Line graphs showing absolute transcript levels of canonical MBF targets (*CLB5* and *RNR1*) and SBF targets (*CLN2* and *PCL2*) in the indicated datasets. Cells were synchronized in early G1 and released into YEPD medium at 37°C. Transcript levels were measured by microarray. (B) Line graphs showing absolute transcript levels of Hcm1 targets in experiments described in (A).

**Figure S3.**
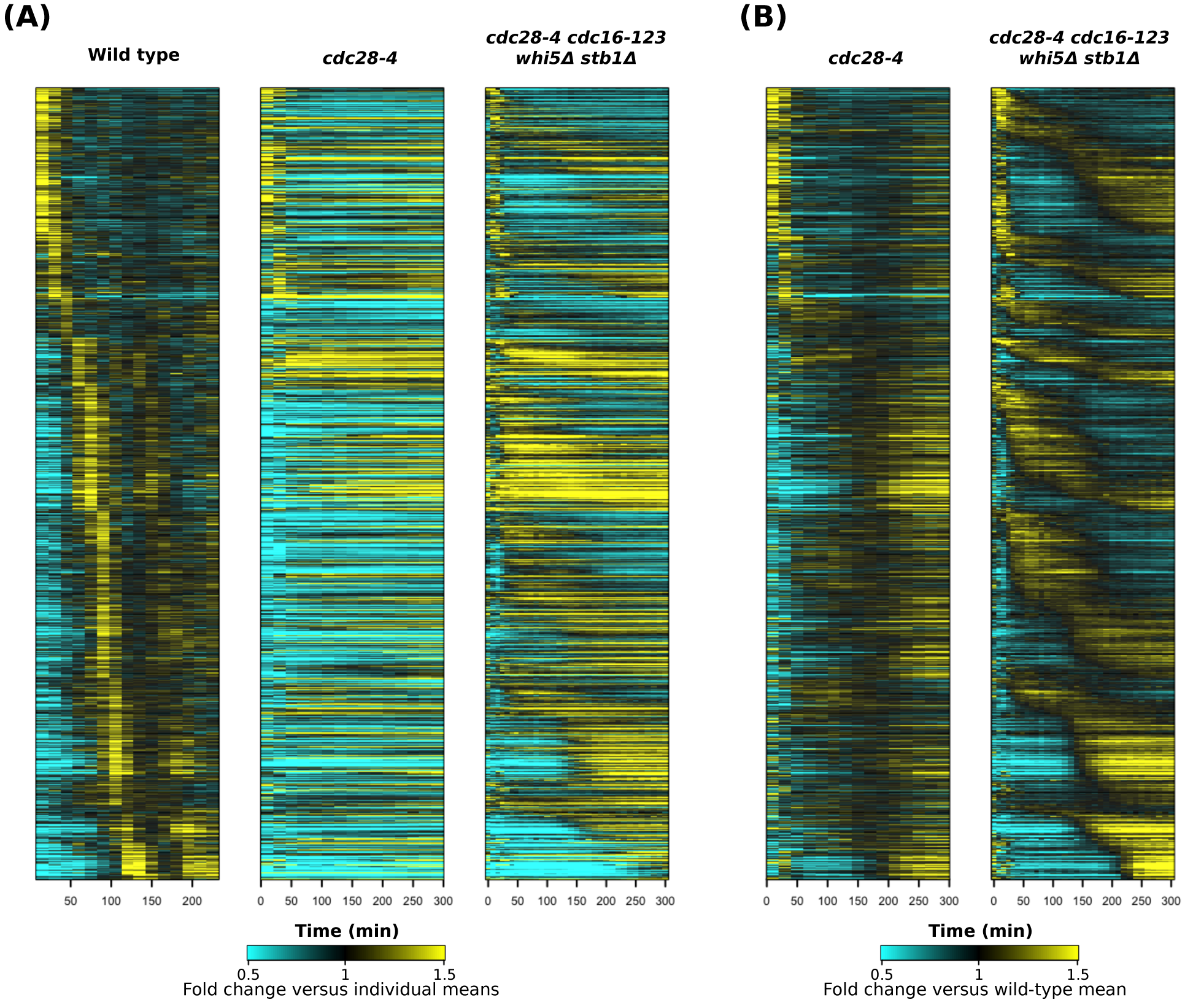
Additional analyses related to the *cdc16-123 cdc28-4 whi5Δ stb1Δ* mutant (related to Figure 4). Heat maps showing transcript dynamics of the same genes from Figure 4A in the same order in indicated strains. Cells were synchronized in G1 by elutriation or α-factor and released into YEPD medium at 37°C. Transcript levels are depicted as fold change relative to mean in individual datasets (A) or relative to wild-type mean (B) (Simmons Kovacs et al., 2012).

**Figure S4.**
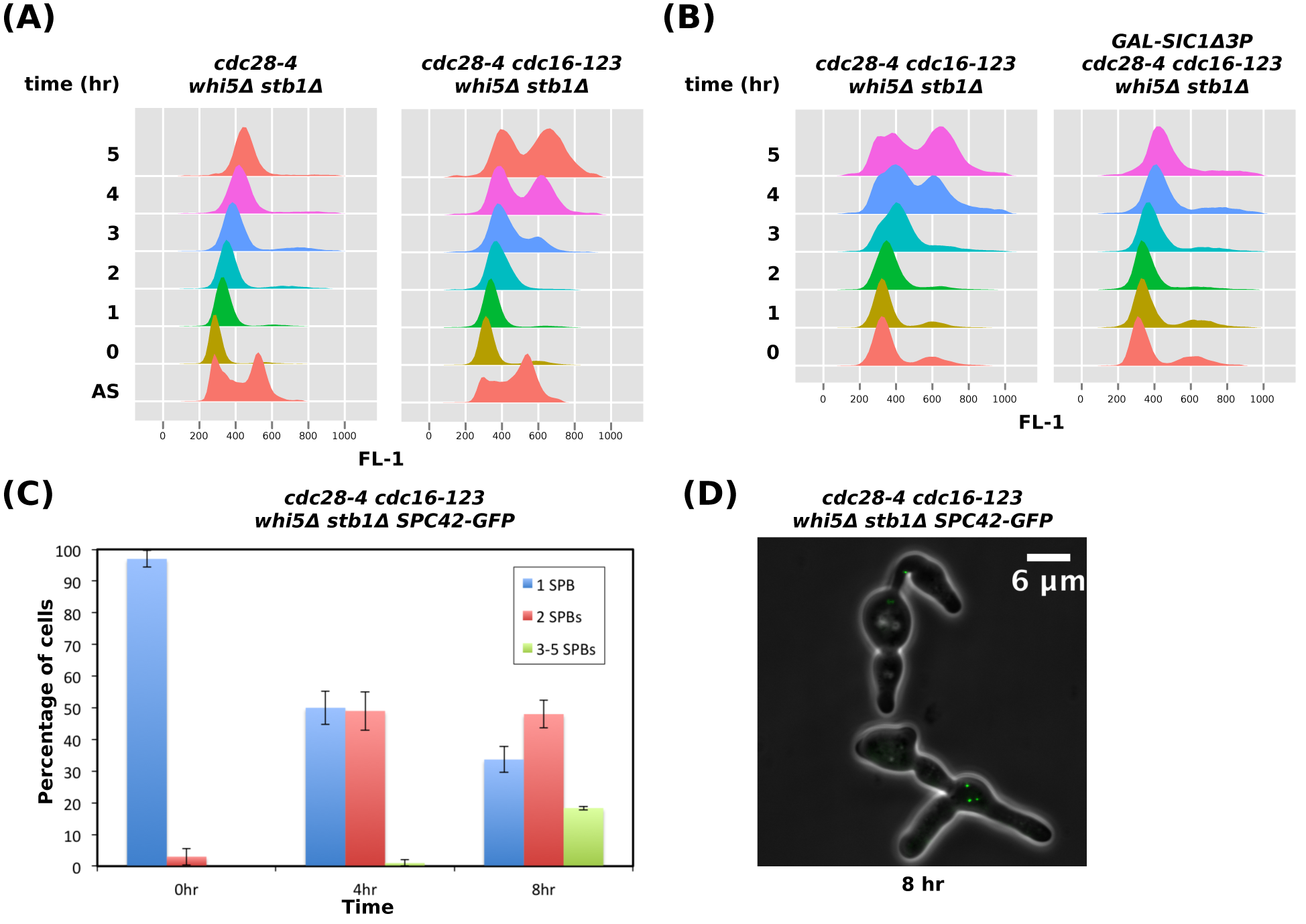
Cell-cycle characterization of the *cdc16-123 cdc28-4 whi5Δ stb1Δ* mutant (related to Figure 3 and Figure 4). (A) DNA content of the indicated strains analyzed by flow cytometry using SYTOX Green during the time course. Cells were synchronized by α-factor in YEPG at 25°C and released into YEPD medium at 37°C. (B) DNA content of the indicated strains analyzed by flow cytometry using SYTOX Green during the time course. Cells were synchronized by α-factor in YEP-sucrose at 25°C, continued the α-factor arrest in YEPG for 40 minutes to induce the overexpression of hyperstable Clb-CDK inhibitor SicΔ3P, and then released into YEPD medium at 37°C. (C) Bar plot showing percentages of cells with different numbers of spindle pole bodies (SPBs) in the *cdc16-123 cdc28-4 whi5Δ stb1Δ* mutant at indicated time points. Cells carrying endogenously tagged *SPC42-GFP* (a marker of spindle pole bodies) were synchronized in G1 by α-factor in YEPG medium, released into YEPD at 37°C, and fixed at indicated time points for counting by fluorescent microscopy. Mean ± s.d. from three independent experiments is shown. (D) Microscopic images of the *cdc16-123 cdc28-4 whi5Δ stb1Δ SPC42-GFP* mutant from experiments described in (C).

**Figure S5.**
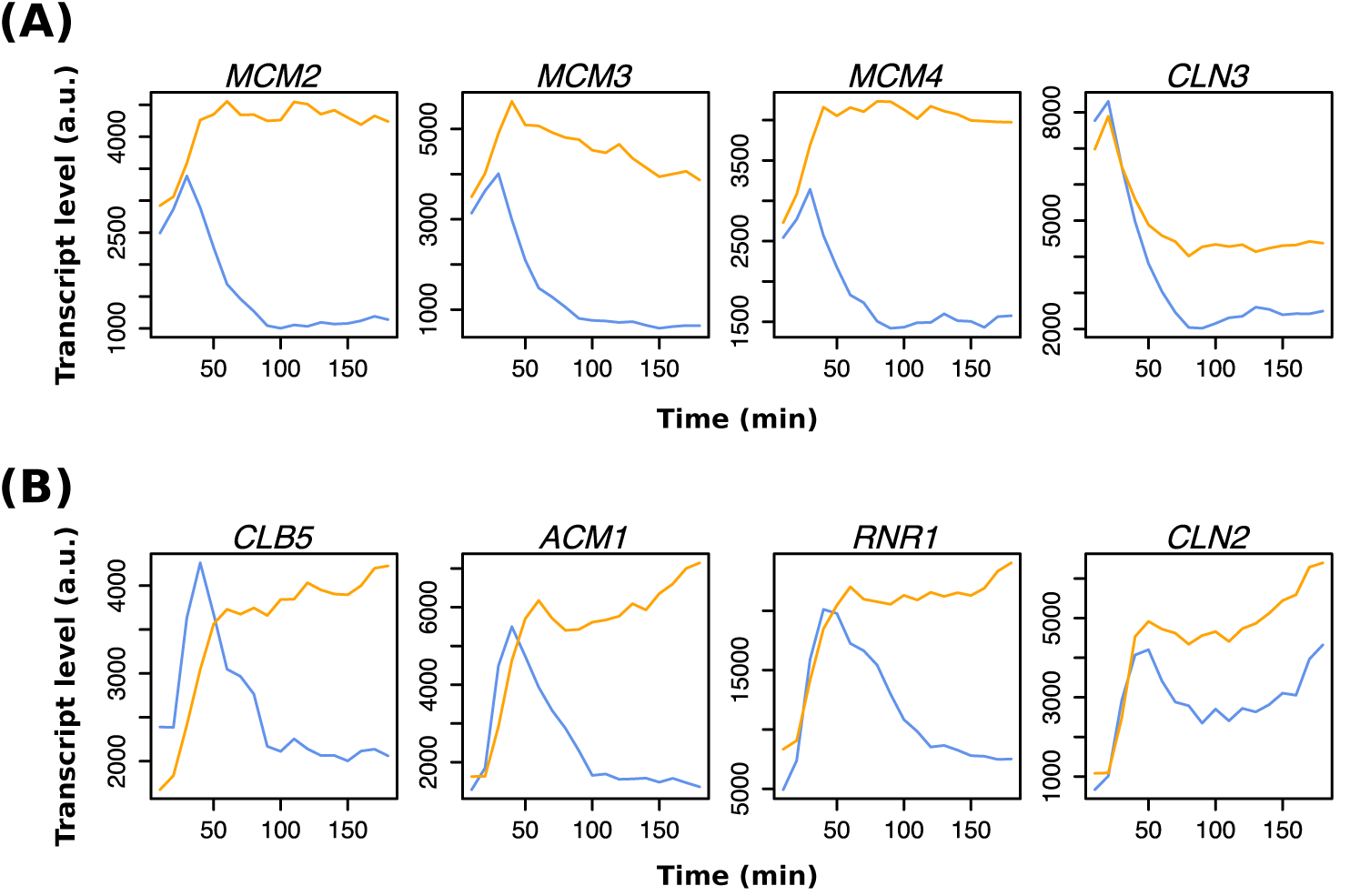
Additional analyses related to the *GAL-SICΔ3P* cells (related to Figure 5). (A) Line graphs showing absolute transcript levels of canonical Mcm1 targets in the *GAL-SIC1Δ3P* cells (blue) or the *GAL-SIC1Δ3P nrm1Δ yhp1Δ yox1Δ* cells (yellow). Cells were synchronized in early G1 and released into YEPD medium at 37°C. Transcript levels were measured by microarray. (B) Line graphs showing absolute transcript levels of SBF/MBF targets in experiments described in (A).

**Table S3.** Strains used in this study

| **Table S3 Strains used in this study** |  |  |
| --- | --- | --- |
| **Strain** | **Genotype** | **Source** |
| SBY2457 | *MATa BAR1 cdc28-4 CDC3-mCherry::LEU2* | This study |
| SBY2459 | *MATa bar1 cdc28-4 whi5Δ::hphMX4 stb1Δ::KanMX6 CDC3-mCherry::LEU2* | This study |
| SBY2356 | *MATa BAR1 cdc28-4 whi5Δ::hphMX4 stb1Δ::KanMX6* | This study |
| SBY2424 | *MATa bar1 cdc28-4 whi5Δ::hphMX4 stb1::KanMX6* | This study |
| SBY2435 | *MATa bar1 YHP1-13myc::KanMX6* | This study |
| SBY2436 | *MATa bar1 cdc28-4 whi5Δ::hphMX4 stb1Δ::KanMX6 YHP1-13myc::TRP1* | This study |
| SBY2398 | *MATa bar1 cdc16-123 cdc28-4 whi5Δ::hphMX4 stb1Δ::KanMX6 YHP1-13myc::TRP1* | This study |
| SBY2437 | *MATa bar1 NRM1-13myc::TRP1* | This study |
| SBY2442 | *MATa bar1 cdc28-4 whi5Δ::hphMX4 stb1Δ::KanMX6 NRM1-13myc::TRP1* | This study |
| SBY2444 | *MATa bar1 cdc16-123 cdc28-4 whi5Δ::hphMX4 stb1Δ::KanMX6 NRM1-13myc::TRP1* | This study |
| SBY2467 | *MATa bar1 NDD1-13myc::TRP1* | This study |
| SBY2446 | *MATa bar1 cdc28-4 whi5Δ::hphMX4 stb1Δ::KanMX6 NDD1-13myc::TRP1* | This study |
| SBY2448 | *MATa bar1 cdc16-123 cdc28-4 whi5Δ::hphMX4 stb1Δ::KanMX6 NDD1-13myc::TRP1* | This study |
| SBY2395 | *MATa bar1 cdc16-123 cdc28-4 whi5Δ::hphMX4 stb1::KanMX6* | This study |
| SBY2451 | *MATa bar1 cdc16-123 cdc28-4 whi5Δ::hphMX4 stb1Δ::KanMX6 GAL-SIC1Δ3P::HIS2* | This study |
| SBY2508 | *MATa bar1 cdc16-123 cdc28-4 whi5Δ::hphMX4 stb1Δ::KanMX6 SPC42-GFP::TRP1* | This study |
| SBY1399 | *MATa bar1 GAL-SIC1Δ3P::URA3* | (Simmons Kovacs et al., 2012) |
| SBY2320 | *MATa bar1 GAL-SIC1Δ3P::URA3 nrm1Δ::natMX4 yhp1Δ::hphMX4 yox1Δ::kanMX6* | This study |

